# Identification of viruses with the potential to infect human

**DOI:** 10.1101/597963

**Authors:** Zheng Zhang, Zena Cai, Zhiying Tan, Congyu Lu, Gaihua Zhang, Yousong Peng

## Abstract

The virus has caused much mortality and morbidity to humans, and still posed a serious threat to the global public health. The virome with the human-infection potential is far from complete. Novel viruses have been discovered at an unprecedented pace as the rapid development of viral metagenomics. However, there is still a lack of a method for rapidly identifying the virus with the human-infection potential. This study built several machine learning models for discriminating the human-infecting viruses from other viruses based on the frequency of k-mers in the viral genomic sequences. The k-nearest neighbor (KNN) model could predict the human-infecting virus with an accuracy of over 90%. Even for the KNN models built on the contigs as short as 1kb, they performed comparably to those built on the viral genomes, suggesting that the models could be used to identify the human-infecting virus from the viral metagenomic sequences. This work could help for discovery of novel human-infecting virus in metagenomics studies.

## Introduction

Viruses are the most abundant biological entities on Earth and exist in all habitats of the world (Paez-Espino, Eloe-Fadrosh et al. 2016). They can infect all kinds of organisms ranging from the bacteria to animals, including the human. Humans are constantly exposed to a large diversity of viruses, and over two-thirds of all human pathogens belong to viruses (Woolhouse and Gaunt 2007). The viruses have caused huge mortality and morbidity to the human society in human history, such as the devastating smallpox and Spanish flu. Although much progress have been made in prevention and control of viral infectious disease, recent serial outbreaks caused by the Middle East Respiratory Syndrome coronavirus (Breban, Riou et al. 2013), avian influenza H7N9 virus (Gao, Cao et al. 2013), Ebola virus (Maganga, Kapetshi et al. 2014) and Zika virus (Mlakar, Korva et al. 2016) indicate that the viruses still pose a serious threat to the global public health.

The virome which could infect humans is far from complete. Generally speaking, a new pathogen would not be identified until it caused epidemics or pandemics. Many viruses may remain undiscovered, even if they have already entered into human populations (Rosenberg 2015). Traditional diagnostic methods such as polymerase chain reaction, immunological assays and pan-viral microarrays are limited in their ability to identify novel viruses with the potential to infect humans. Rapid development of viral metagenomic sequencing in recent years provides a powerful high-throughput and culture-independent means to identify new viruses, which leads to accumulation of new viruses at an unprecedented pace (Alavandi and Poornima 2012). The Global Virome Project (GVP), which was proposed and initiated in the beginning of 2018, estimated that there were over 1.67 million yet-to-be-discovered viruses in animal reservoirs, and between 631,000 to 827,000 of these unknown viruses have the capacity to infect humans (Carroll, Daszak et al. 2018). Therefore, it is in great need to develop rapid methods for identifying viruses with the potential to infect humans.

Two kinds of methods, the sequence alignment-based and alignment-free methods, have been developed to predict the host for viruses. For example, several methods based on k-mers, which were extracted from viral genomes (Ahlgren, Ren et al. 2016, Li and Sun 2018), and several methods based on sequence blast, have been developed to predict the host of the phage (Bolotin, Quinquis et al. 2005, Edwards, McNair et al. 2015). Some studies have also attempted to identify the human virus with these methods. For example, Xu et al. (Xu, Tan et al. 2017) developed SVM models to predict the host of influenza viruses based on word vectors. However, all these studies have focused some type of viruses, such as coronavirus and influenza virus. The methods they developed were not suitable for identification of novel human-infecting viruses from the viral metagenomic sequences. Here, we attempted to build machine learning models for rapid identification of viruses with the potential to infect human.

## Materials and Methods

### Virus-host interactions and viral genomes

The virus with specific hosts and the viral genomic sequences were obtained from the database of Virus-Host DB (available at https://www.genome.jp/virushostdb/) on July 15, 2018 (Mihara, Nishimura et al. 2016). After removing the viroid and satellites, and the viruses with genomic sequence less than 1kb, a total of 9428 viruses were kept for further analysis, which included 1236 viruses infecting humans, which were defined as human-infecting viruses, and 8192 viruses infecting other species.

### Machine learning models

The machine learning models of k-nearest neighbor (KNN) (k=1), support vector machine (SVM) (using the linear kernel function), gaussian naive bayes classifier (GNBC), random forest (RF) (with default settings) and logistic regression (LR) (with default settings) were built with the default parameters using the package “scikit-learn” (version 0.20.2) (Pedregosa, Varoquaux et al. 2011) in Python (version 3.6.2). Because the number of human-infecting viruses was much smaller than that of other viruses, the functions of “BalanceBaggingClassifier” (Breiman 1996, Barandiaran 1998) and “BalanceRandomForest” (Chen, Liaw et al. 2004) in the package of “imbalanced-learn” (version 0.4.3) in Python were used to deal with the imbalance between the human-infecting virus and other viruses in the modeling, with the parameter of “n_estimators” set to be 100.

Ten-fold cross-validations were used to evaluate the predictive performances of the machine learning models, and were conducted using the functions of ‘StratifiedKFold’ in the package “scikit-learn” in Python. To measure the predictive performances of the machine learning models, the area under the receiver operating characteristics (ROC) curve (AUC), the accuracy, recall rate, specificity and predictive precision were calculated for each model.

### Statistical analysis

All the statistical analysis was conducted in R (version 3.5.0).

## Results

### Taxonomy distribution of viruses

A total of 9428 viruses were used in the study. Among them, 1236 viruses were reported to infect humans, which were defined as the human-infecting virus. They included 1043 viruses exclusively infecting humans and 193 viruses both infecting humans and other species. The human-infecting virus covered all groups of viruses in the Baltimore classification (Figure 1A). Over 60% of the human-infecting virus belonged to single-stranded RNA (ssRNA) viruses, which also had the highest ratio of human-infecting viruses (28%). The second largest component (25%) of the human-infecting virus was the double-stranded DNA (dsDNA) virus. On the level of family, the human-infecting virus comprised of viruses from 30 viral families. Among them, the families of Caliciviridae (ssRNA), Picornaviridae (ssRNA) and Papillomaviridae (dsDNA) were the three most abundant ones, which accounted for nearly 60% of the human-infecting virus.

**Figure 1.**
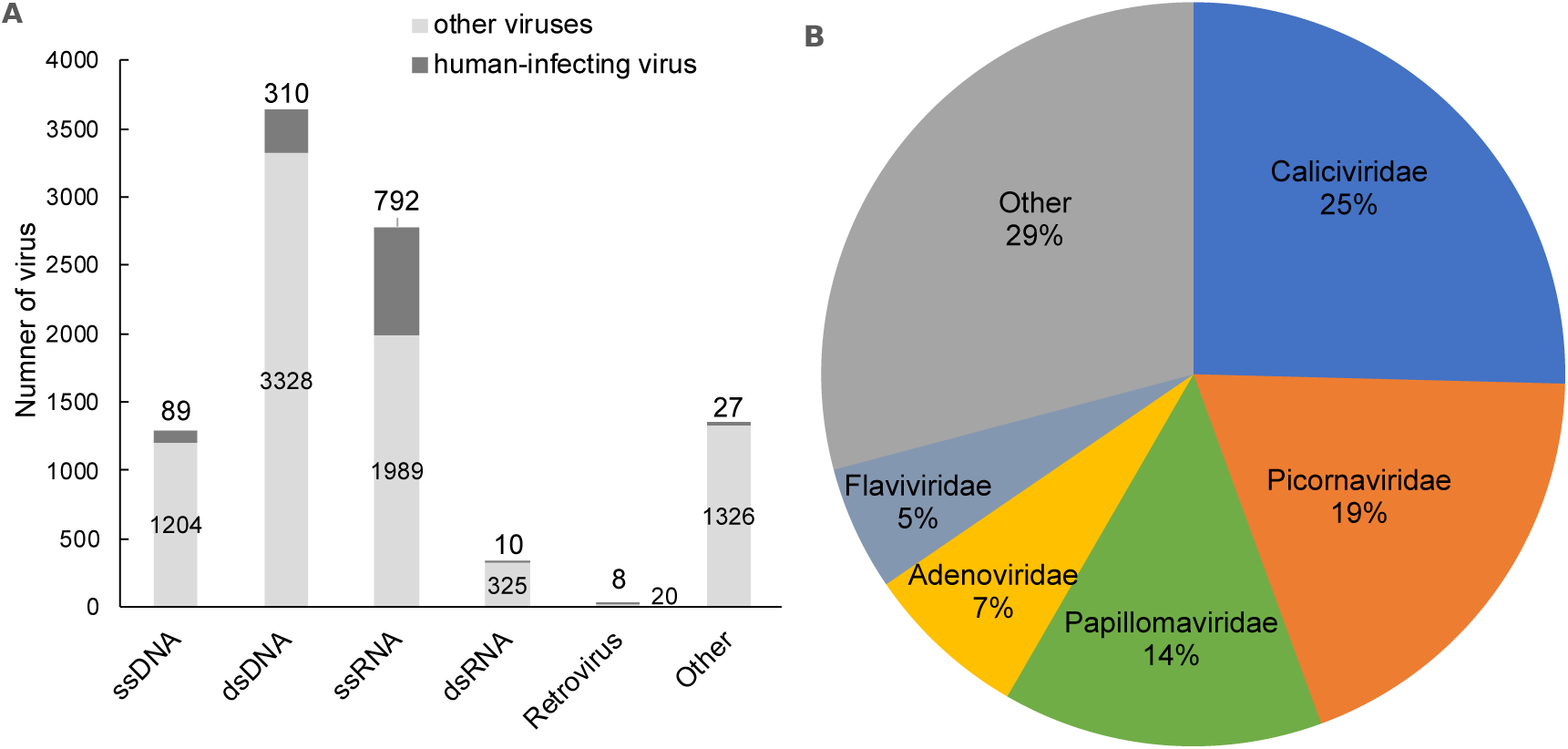
Analysis of the taxonomy distribution of all viruses used. (A) Taxonomy distribution of the human-infecting and other viruses in the Baltimore classification. (B) Taxonomy distribution of the human-infecting virus by viral family.

Besides the human-infecting virus, there were a total of 8192 viruses which infect other species except the human, including the archaea, bacteria, fungi, plant, animal, and so on. They also covered all groups of viruses in the Baltimore classification (Figure 1A), and comprised of viruses from 91 viral families.

### Machine-learning models for identifying the human-infecting virus based on k-mer frequencies in the genome

To discriminating the human-infecting virus from other viruses, several machine-learning models, including k-nearest neighbor (KNN), random forest (RF), gaussian naive bayes classifier (GNBC), support-vector machine (SVM) and logistic regression (LR), were built based on k-mer frequencies in the viral genome. K-mers of one to six nucleotides were used in the models to evaluating the influence of the k-mer length on the model performance (Table S1). Figure 2 showed that the AUC of GNBC, SVM and LR models increased as the increase of the k-mer length from one to six. While for KNN and RF, the AUCs of them peaked at k-mer length of 4 and 3, respectively; then, they began to decrease as the increase of k-mer length. The AUCs of KNN and RF were much larger than those of other models, including GNBC, SVM and LR, at all length of k-mers, suggesting that the KNN and RF outperformed other models in discriminating the human-infecting virus from other viruses. The best performance was achieved on the KNN model with the k-mers of four nucleotides. The model not only had the best overall performance, with accuracy of 0.90 and AUC of 0.92, respectively, but also had the best ability of capturing the human-infecting virus, with recall rate of 0.94.

**Figure 2.**
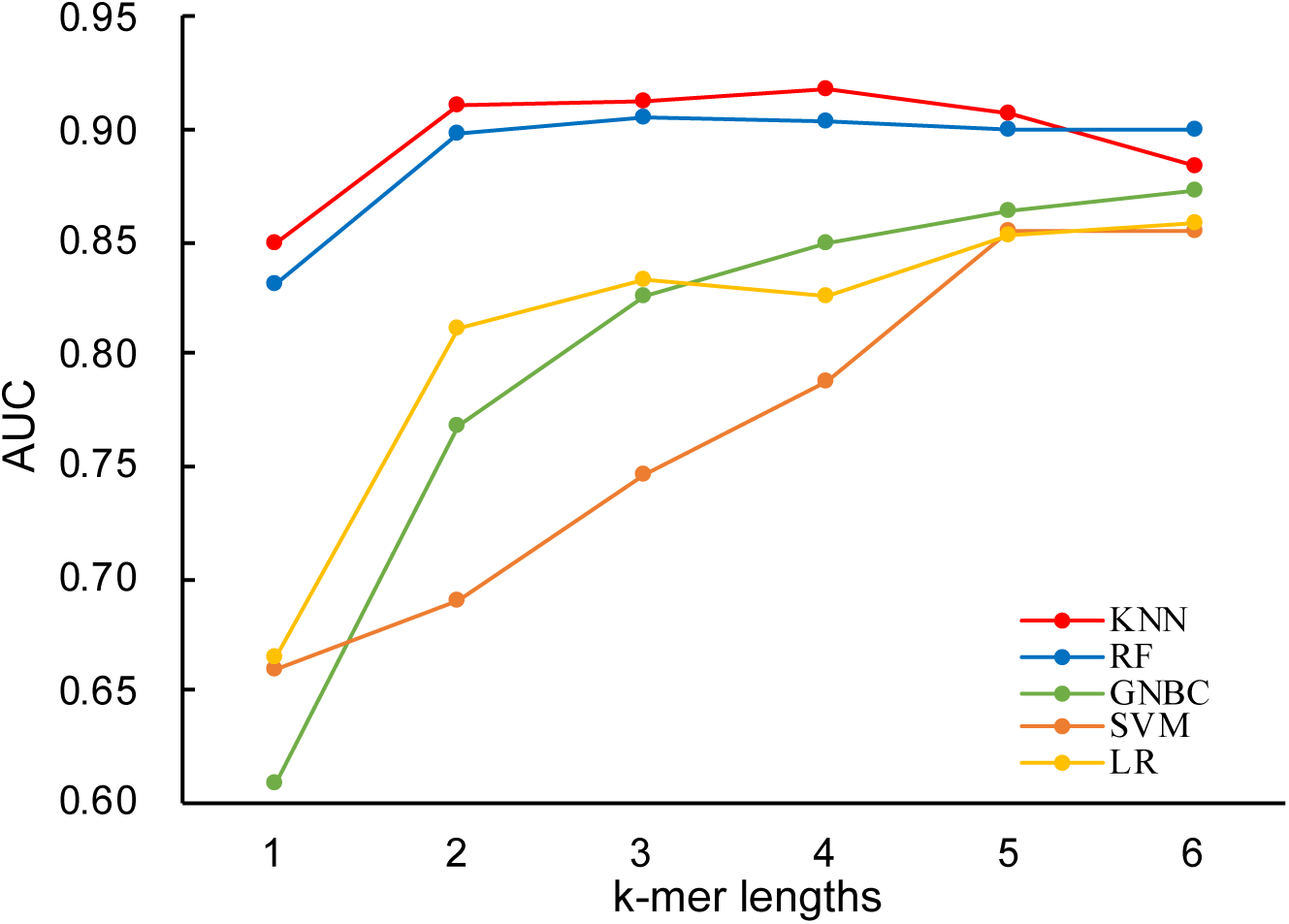
The AUCs for machine learning models with k-mer length ranging from one to six.

### Identifying the human-infecting virus based on contigs of varying length

In the metagenomics studies, contigs of varying length were assembled, which range from several hundreds to several thousands of nucleotides. Rapid determination of the human-infecting virus from metagenomic sequences is important for early warnings of newly emerging viruses. To mimic the metagenomic sequences, viral genomes were split into non-overlapping contigs of 500, 1,000, 3,000, 5,000 and 10,000 nucleotides long. The number of contigs of varying length in the human-infecting virus and other viruses were listed in Table S2.

Since the KNN model performed best in discriminating the human-infecting virus from other viruses, it was used to identify the human-infecting virus based on contigs of varying length. For the contigs of a given length, KNN models for discriminating the human-infecting virus and other viruses were built with k-mers of different size (Table S3). As was shown in Figure 3, for contigs of all length, the AUC of KNN models increased as the increase of k-mer length from one to four. The best performance was achieved for all models with k-mers of four nucleotides long. The longer the contigs were, the better the models performed in identifying the human-infecting virus (Figure 3 and Table 2). The model built on the contigs of 10,000 nucleotides had an AUC of 0.96 and a recall rate of 0.99, suggesting the model could capture most human-infecting viruses. While on the contig of 500 nucleotides, the model only had an AUC of 0.86 and a recall rate of 0.88. Attention to note, for models built on the contigs of 1000 nucleotides or longer, they performed similarly to those built on the viral genome.

**Table 1.**
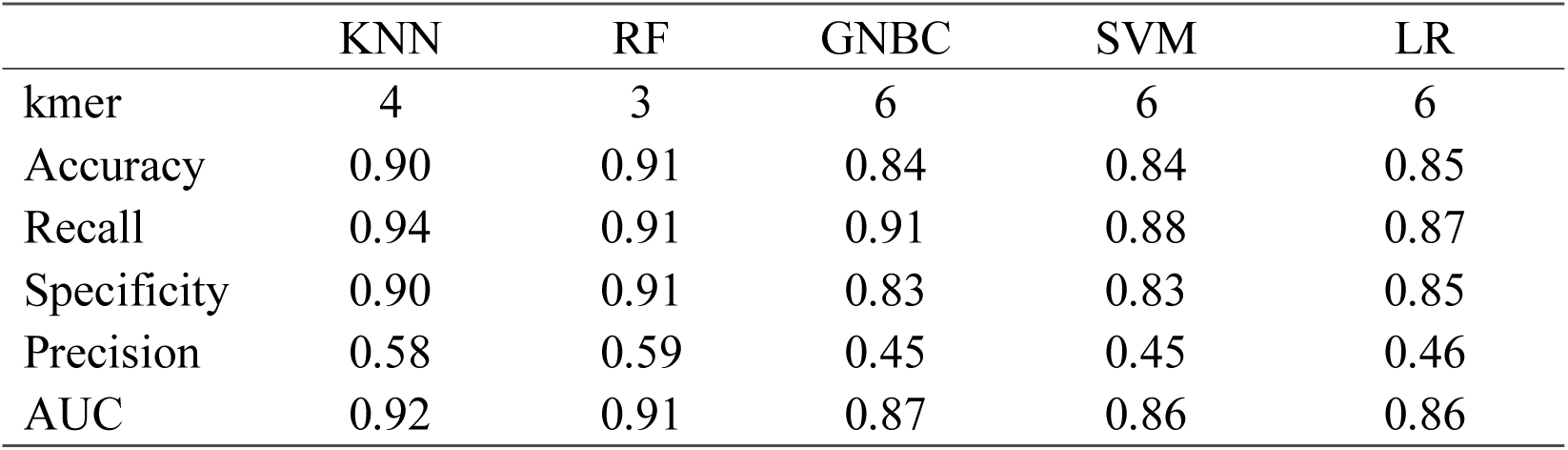
Best performance of each kind of machine learning model.

|  | KNN | RF | GNBC | SVM | LR |
| --- | --- | --- | --- | --- | --- |
| kmer | 4 | 3 | 6 | 6 | 6 |
| Accuracy | 0.90 | 0.91 | 0.84 | 0.84 | 0.85 |
| Recall | 0.94 | 0.91 | 0.91 | 0.88 | 0.87 |
| Specificity | 0.90 | 0.91 | 0.83 | 0.83 | 0.85 |
| Precision | 0.58 | 0.59 | 0.45 | 0.45 | 0.46 |
| AUC | 0.92 | 0.91 | 0.87 | 0.86 | 0.86 |

**Table 2.**
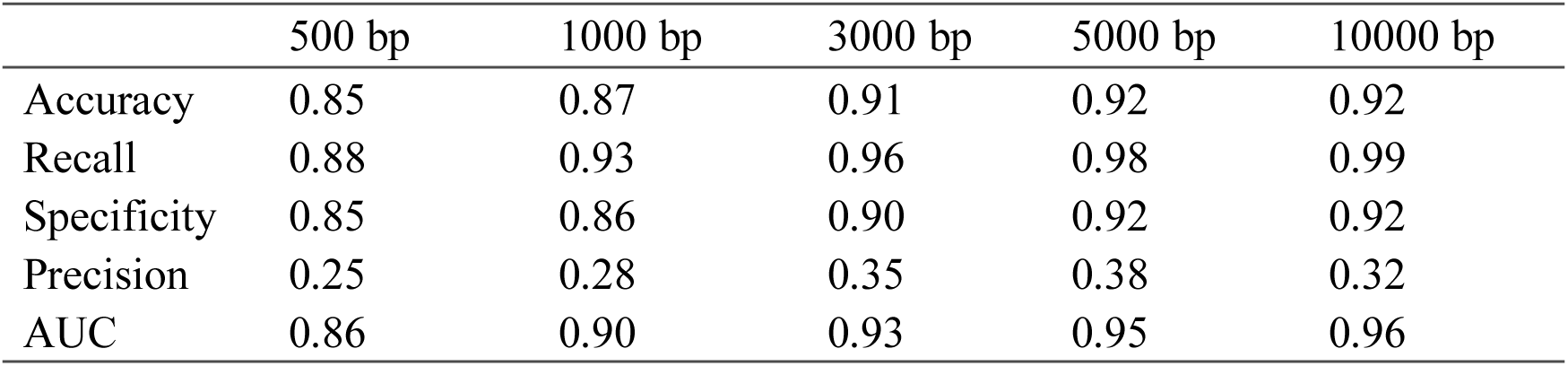
Best performance of KNN models built on contigs of varying length with k-mers of four nucleotides used in modeling.

**Figure 3.**
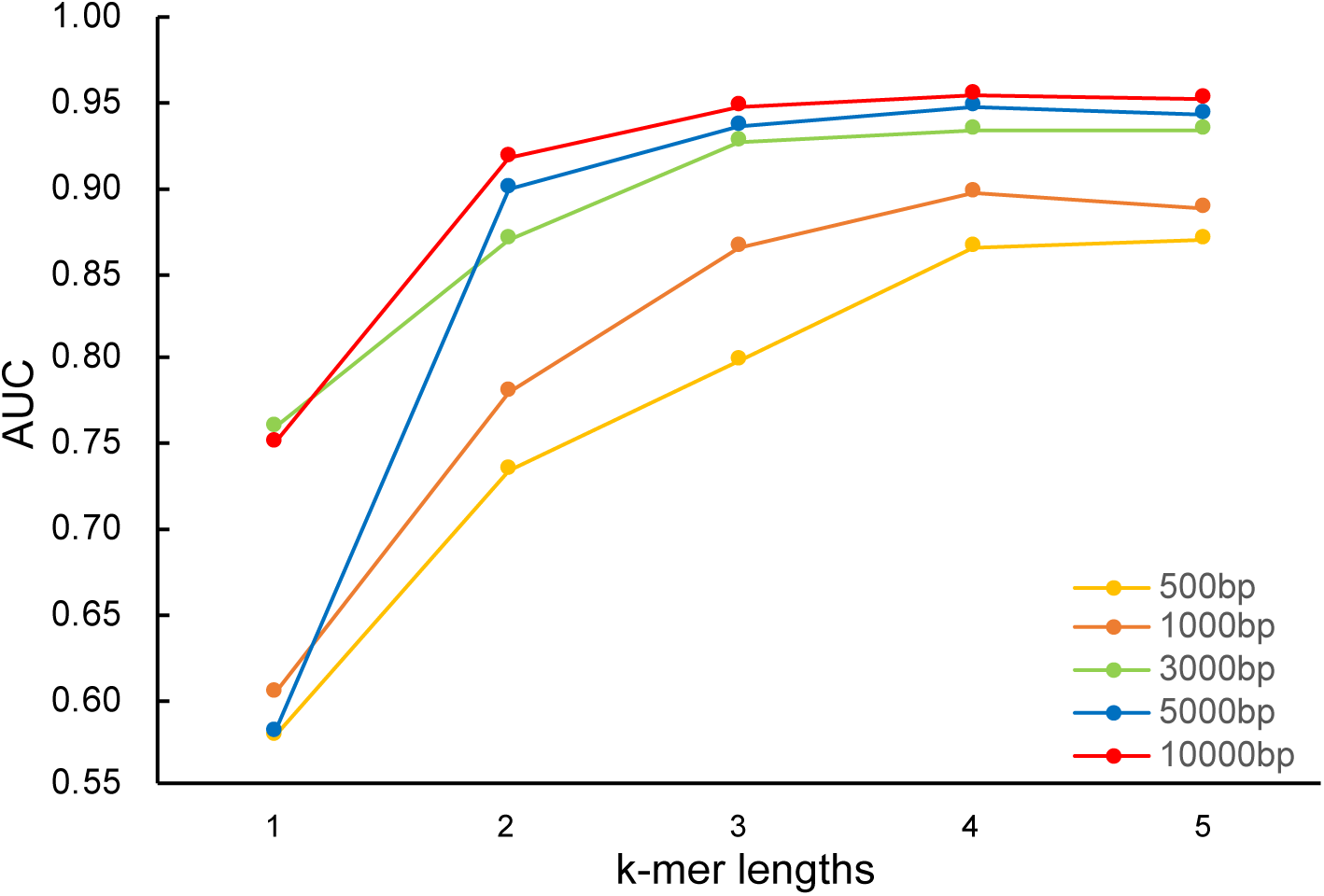
The AUCs for KNN models built on contigs of varying length, with k-mer length ranging from one to five.

## Discussion

Identification of human-infecting viruses is important for early warnings of newly emerging viruses. As the rapid accumulation of viral metagenomic sequences, more and more novel viruses have been discovered. Unfortunately, there is currently a lack of rapid methods for identification of viruses with the potential to infect humans. As far as we know, this study for the first time attempted to discriminate the human-infecting virus from non-human-infecting virus in the perspective of virome. The KNN model could predict the human-infecting virus with high accuracy and sensitivity. Even for the KNN models built on the contigs as short as 1kb, they performed comparably to those built on the viral genomes. This suggested that the models built here could be used in metagenomics for identifying the viruses with the potential of infecting humans.

The human-infecting virus used here has different ability of infecting humans. Some of them are human viruses and have been circulating in human populations for a long time, such as the human papillomavirus (Baseman and Koutsky 2005) and yellow fever virus (Akondy, Monson et al. 2009); while some of them were zoonotic and only infect humans rarely, such as the avian influenza virus and coronavirus. As was shown in numerous studies, the zoonotic viruses are very likely to cause epidemics or even pandemics after adaptive mutations. The most recent example is the Zika virus. It rarely caused wide-spread epidemics in humans, even in highly enzootic areas, in the 20^th^ century. However, a few mutations on the viral genome facilitated its rapid spread in human populations. The virus has caused epidemics in more than 50 countries and has infected more than one million people since 2014 (Prevention and Control 2016, de Oliveira Garcia 2019). Taken together, it is difficult to distinguish between viruses with different extent of infecting humans. Therefore, the human-infecting virus with varying ability of infecting humans were considered equally in the modeling.

There are some limitations to this study. Firstly, the human-infecting virus is far from complete when compared to that estimated by the GVP. But they covered viruses of 30 families in all groups of the Baltimore classification, which was similar to that estimated by the GVP. Besides, the machine learning models had excellent performances in discriminating the human-infecting virus from other viruses. They could help much in discovering novel human-infecting viruses. infect humans after adaptive mutations. Secondly, the number of human-infecting virus is much smaller than that of other viruses. Such a large imbalance may hinder accurate modeling. Here, the under-sampling method was used to deal with the imbalance problem. The KNN model not only achieved excellent overall performances, but also had high sensitivity in identifying the human-infecting virus.

In conclusion, this study for the first time built the computational models to predict the human-infecting virus in the perspective of virome. The high accuracy and sensitivity of the KNN model built on the viral contigs of varying length suggests that the model could be used to identify the human-infecting virus from the viral metagenomic sequences. This work could help for discovery of novel human-infecting virus in metagenomics studies.

## Reference

Ahlgren, N. A., et al. (2016). “Alignment-free oligonucleotide frequency dissimilarity measure improves prediction of hosts from metagenomically-derived viral sequences.” Nucleic acids research 45(1): 39–53.

Akondy, R. S., et al. (2009). “The yellow fever virus vaccine induces a broad and polyfunctional human memory CD8+ T cell response.” The Journal of Immunology 183(12): 7919–7930.

Alavandi, S. and M. Poornima (2012). “Viral metagenomics: a tool for virus discovery and diversity in aquaculture.” Indian Journal of Virology 23(2): 88–98.

Barandiaran, I. (1998). “The random subspace method for constructing decision forests.” IEEE Trans. Pattern Anal. Mach. Intell 20(8).

Baseman, J. G. and L. A. Koutsky (2005). “The epidemiology of human papillomavirus infections.” Journal of clinical virology 32: 16–24.

Bolotin, A., et al. (2005). “Clustered regularly interspaced short palindrome repeats (CRISPRs) have spacers of extrachromosomal origin.” Microbiology 151(8): 2551–2561.

Breban, R., et al. (2013). “Interhuman transmissibility of Middle East respiratory syndrome coronavirus: estimation of pandemic risk.” The Lancet 382(9893): 694-699.

Breiman, L. (1996). “Bagging predictors.” Machine learning 24(2): 123–140.

Carroll, D., et al. (2018). “The global virome project.” Science 359(6378): 872-874.

Chen, C., et al. (2004). “Using random forest to learn imbalanced data.” University of California, Berkeley 110: 1–12.

de Oliveira Garcia M. H. (2019). “Zika: the continuing threat.” Bull World Health Organ 97: 6–7.

Edwards, R. A., et al. (2015). “Computational approaches to predict bacteriophage–host relationships.” FEMS microbiology reviews 40(2): 258–272.

Gao, R., et al. (2013). “Human infection with a novel avian-origin influenza A (H7N9) virus.” New England Journal of Medicine 368(20): 1888–1897.

Li, H. and F. Sun (2018). “Comparative studies of alignment, alignment-free and SVM based approaches for predicting the hosts of viruses based on viral sequences.” Scientific reports 8(1): 10032.

Maganga, G. D., et al. (2014). “Ebola virus disease in the Democratic Republic of Congo.” New England Journal of Medicine 371(22): 2083–2091.

Mihara, T., et al. (2016). “Linking virus genomes with host taxonomy.” Viruses 8(3): 66.

Mlakar, J., et al. (2016). “Zika virus associated with microcephaly.” New England Journal of Medicine 374(10): 951–958.

Paez-Espino, D., et al. (2016). “Uncovering Earth’s virome.” Nature 536(7617): 425.

Pedregosa, F., et al. (2011). “Scikit-learn: Machine learning in Python.” Journal of machine learning research 12(Oct): 2825–2830.

Prevention, E. C. f. D. and Control (2016). Zika virus epidemic in the Americas: potential association with microcephaly and Guillain-Barré syndrome (first update), ECDC Stockholm.

Rosenberg, R. (2015). “Detecting the emergence of novel, zoonotic viruses pathogenic to humans.” Cellular and molecular life sciences 72(6): 1115–1125.

Woolhouse, M. and E. Gaunt (2007). “Ecological origins of novel human pathogens.” Critical reviews in microbiology 33(4): 231–242.

Xu, B., et al. (2017). “Predicting the host of influenza viruses based on the word vector.” PeerJ 5: e3579.

