## Supplemental Tables for "Identification of viruses with the potential to infect human"

**Supplementary Tables**

**Table S1.** The performance of machine learning models with k-mer length ranging from one to six.

|  | KNN | RF | GNBC | SVM | LR |
| --- | --- | --- | --- | --- | --- |
| kmer=1 | |  |  |  |  |
| accuracy | 0.85 | 0.84 | 0.45 | 0.59 | 0.61 |
| recall | 0.85 | 0.82 | 0.83 | 0.77 | 0.74 |
| specificity | 0.85 | 0.85 | 0.39 | 0.56 | 0.59 |
| precision | 0.46 | 0.44 | 0.17 | 0.21 | 0.22 |
| AUC | 0.85 | 0.83 | 0.61 | 0.66 | 0.67 |
| kmer=2 | |  |  |  |  |
| accuracy | 0.90 | 0.90 | 0.74 | 0.49 | 0.78 |
| recall | 0.92 | 0.89 | 0.81 | 0.95 | 0.86 |
| specificity | 0.90 | 0.91 | 0.73 | 0.42 | 0.77 |
| precision | 0.58 | 0.59 | 0.31 | 0.20 | 0.36 |
| AUC | 0.91 | 0.90 | 0.77 | 0.69 | 0.81 |
| kmer=3 | |  |  |  |  |
| accuracy | 0.90 | 0.91 | 0.80 | 0.59 | 0.80 |
| recall | 0.93 | 0.91 | 0.86 | 0.95 | 0.88 |
| specificity | 0.90 | 0.91 | 0.79 | 0.54 | 0.79 |
| precision | 0.58 | 0.59 | 0.38 | 0.24 | 0.38 |
| AUC | 0.91 | 0.91 | 0.83 | 0.74 | 0.83 |
| kmer=4 | |  |  |  |  |
| accuracy | 0.90 | 0.90 | 0.82 | 0.68 | 0.81 |
| recall | 0.94 | 0.92 | 0.89 | 0.94 | 0.86 |
| specificity | 0.90 | 0.89 | 0.81 | 0.64 | 0.80 |
| precision | 0.58 | 0.57 | 0.41 | 0.28 | 0.39 |
| AUC | 0.92 | 0.90 | 0.85 | 0.79 | 0.83 |
| kmer=5 | |  |  |  |  |
| accuracy | 0.88 | 0.89 | 0.83 | 0.81 | 0.83 |
| recall | 0.93 | 0.91 | 0.91 | 0.91 | 0.89 |
| specificity | 0.88 | 0.89 | 0.82 | 0.79 | 0.83 |
| precision | 0.53 | 0.56 | 0.43 | 0.40 | 0.44 |
| AUC | 0.90 | 0.90 | 0.86 | 0.85 | 0.86 |
| kmer=6 | |  |  |  |  |
| accuracy | 0.90 | 0.90 | 0.84 | 0.84 | 0.85 |
| recall | 0.86 | 0.90 | 0.91 | 0.88 | 0.87 |
| specificity | 0.90 | 0.90 | 0.83 | 0.83 | 0.85 |
| precision | 0.57 | 0.57 | 0.45 | 0.45 | 0.46 |
| AUC | 0.88 | 0.90 | 0.87 | 0.86 | 0.86 |

**Table S2**. The number of contigs of varying length generated from the virus genomes.

| Contig Length (bp) | human-infecting virus | other viruses | Total |
| --- | --- | --- | --- |
| 500 | 35,843 | 627,587 | 663,430 |
| 1000 | 17,579 | 311,655 | 329,234 |
| 3000 | 5435 | 100,775 | 106,210 |
| 5000 | 3053 | 58931 | 61984 |
| 10000 | 1043 | 27410 | 28453 |

**Table S3.** The performance of KNN models built on contigs of varying length with k-mer length ranging from one to six. ^a^ Not applicable because the kinds of k-mers are much larger than the contig length.

|  | 500 bp | 1000 bp | 3000 bp | 5000 bp | 10000 bp |
| --- | --- | --- | --- | --- | --- |
| kmer=1 |  |  |  |  |  |
| accuracy | 0.62 | 0.66 | 0.79 | 0.62 | 0.79 |
| recall | 0.53 | 0.54 | 0.73 | 0.53 | 0.71 |
| specificity | 0.63 | 0.67 | 0.79 | 0.63 | 0.79 |
| precision | 0.08 | 0.08 | 0.16 | 0.07 | 0.11 |
| AUC | 0.58 | 0.60 | 0.76 | 0.58 | 0.75 |
| kmer=2 |  |  |  |  |  |
| accuracy | 0.77 | 0.81 | 0.87 | 0.89 | 0.91 |
| recall | 0.69 | 0.75 | 0.87 | 0.91 | 0.93 |
| specificity | 0.78 | 0.81 | 0.87 | 0.89 | 0.91 |
| precision | 0.15 | 0.18 | 0.26 | 0.29 | 0.28 |
| AUC | 0.73 | 0.78 | 0.87 | 0.90 | 0.92 |
| kmer=3 |  |  |  |  |  |
| accuracy | 0.81 | 0.85 | 0.90 | 0.91 | 0.92 |
| recall | 0.79 | 0.88 | 0.96 | 0.96 | 0.97 |
| specificity | 0.81 | 0.84 | 0.90 | 0.91 | 0.92 |
| precision | 0.19 | 0.24 | 0.34 | 0.35 | 0.32 |
| AUC | 0.80 | 0.86 | 0.93 | 0.94 | 0.95 |
| kmer=4 |  |  |  |  |  |
| accuracy | 0.85 | 0.87 | 0.91 | 0.92 | 0.92 |
| recall | 0.88 | 0.93 | 0.96 | 0.98 | 0.99 |
| specificity | 0.85 | 0.86 | 0.90 | 0.92 | 0.92 |
| precision | 0.25 | 0.28 | 0.35 | 0.38 | 0.32 |
| AUC | 0.86 | 0.90 | 0.93 | 0.95 | 0.96 |
| kmer=5 |  |  |  |  |  |
| accuracy | 0.94 | 0.84 | 0.91 | 0.92 | 0.92 |
| recall | 0.81 | 0.95 | 0.96 | 0.97 | 0.98 |
| specificity | 0.94 | 0.83 | 0.91 | 0.91 | 0.92 |
| precision | 0.45 | 0.24 | 0.36 | 0.37 | 0.33 |
| AUC | 0.88 | 0.89 | 0.93 | 0.94 | 0.95 |
| kmer=6 | -^a^ | - |  |  |  |
| accuracy | - | - | 0.93 | 0.92 | 0.93 |
| recall | - | - | 0.92 | 0.95 | 0.97 |
| specificity | - | - | 0.93 | 0.92 | 0.93 |
| precision | - | - | 0.42 | 0.39 | 0.33 |
| AUC | - | - | 0.93 | 0.94 | 0.95 |
